# *In silico* identification of metabolic enzyme drug targets in *Burkholderia pseudomallei*

**DOI:** 10.1101/034306

**Authors:** Jean F. Challacombe

## Abstract

The intracellular pathogen *Burkholderia pseudomallei,* which is endemic to parts of southeast Asia and northern Australia, causes the disease melioidosis. Although acute infections can be treated with antibiotics, melioidosis is difficult to cure, and some patients develop chronic infections or a recrudescence of the disease months or years after treatment of the initial infection. *B. pseudomallei* strains have a high level of natural resistance to a variety of antibiotics, and with limited options for new antibiotics on the horizon, new alternatives are needed. The aim of the present study was to characterize the metabolic capabilities of *B. pseudomallei*, identify metabolites crucial for pathogen survival, understand the metabolic interactions that occur between pathogen and host cells, and determine if metabolic enzymes produced by the pathogen might be potential antibacterial targets. This aim was accomplished through genome scale metabolic modeling under different external conditions: 1) including all nutrients that could be consumed by the model, and 2) providing only the nutrients available in culture media. Using this approach, candidate chokepoint enzymes were identified, then knocked out *in silico* under the different nutrient conditions. The effect of each knockout on the metabolic network was examined. When five of the candidate chokepoints were knocked out *in silico*, the flux through the *B. pseudomallei* network was decreased, depending on the nutrient conditions. These results demonstrate the utility of genome-scale metabolic modeling methods for drug target identification in *B. pseudomallei.*

## 1. Introduction

The intracellular pathogen *Burkholderia pseudomallei,* which causes the disease melioidosis, is acquired from the environment in parts of southeast Asia and northern Australia [1,2]. Although acute infections can be treated with antibiotics, melioidosis is difficult to cure, requiring lengthy treatment in two phases for a duration of ~20 weeks [3,4]. Despite antibiotic therapy, some patients have persistent cases that develop into chronic infections, and others experience a recrudescence of the disease months or years after treatment of the initial infection with antibiotics [5]. *B. pseudomallei* strains have a high level of natural resistance to a variety of antibiotics [6–8], and with limited options for new antibiotics on the horizon, alternatives are desperately needed.

The availability of many *B. pseudomallei* genomes and advances in computational analysis methods make possible the rapid identification of novel antibacterial targets by selecting the most likely targets from complete sets of protein coding genes. Previous studies have demonstrated that essential genes present in pathogen genomes, but not in the host, make the best therapeutic targets [9]. Many known antibacterial compounds are enzyme inhibitors [10,11], so metabolic enzymes specific to pathogenic bacteria represent promising drug targets [9].

Enzyme targets in key metabolic pathways have been identified in *B. pseudomallei* and other bacterial pathogens; these pathways include fatty acid biosynthesis [12–14], the glyoxalate shunt [15,16], the chorismate pathway for biosynthesis of aromatic amino acids [17], purine, histidine, 4-aminobenzoate, and lipoate biosynthesis [18,19], leucine, threonine, p-aminobenzoic acid, aromatic compound biosynthesis [20], branched chain amino acid biosynthesis [21], purine metabolism [22]. Other enzyme targets have been identified that are not in pathways - alanine racemase (interconverts L- and D-alanine) [23], superoxide dismutase [24] and cyclic di-GMP phosphodiesterase [25].

Drugs acting on pathogen targets that are not present in the host should not cause significant side effects. However, before the human genome was available, the process of identifying bacterial pathogen-specific drug targets was labor intensive, involving comparison of candidate pathogen targets against all known eukaryotic sequences to filter out targets likely to occur in the human [9]. Since then, various software tools have made the process of *in silico* target identification in pathogen genomes easier. Available *in silico* tools encompass various cheminformatic [26] and bioinformatic [27,28] approaches to identify new protein targets. Among the bioinformatic tools, metabolic pathway/metabolic network analysis has emerged as an efficient *in silico* method to identify candidate metabolic enzyme targets in pathogen genomes.

Several software packages are available to facilitate genome-scale metabolic network analyses [29]. Starting with an annotated pathogen genome, the components of metabolic pathways are identified, curated and refined [30]. The resulting genome-scale metabolic model can be used to integrate omics datasets and to perform various analyses to determine the most likely drug targets [10]. Perhaps the most important task with respect to finding good candidate targets is metabolic chokepoint identification. By definition, a chokepoint enzyme either consumes a unique substrate or produces a unique product in the pathogen metabolic network [31]. Inhibition of chokepoint enzymes may disrupt crucial metabolic processes in the pathogen, so chokepoints that are essential to the pathogen represent good potential drug targets [32–34].

The aim of the present study was to characterize the metabolic capabilities of *B. pseudomallei,* identify metabolites and aspects of the metabolic network crucial for pathogen survival, understand the metabolic interactions that occur between pathogen and host cells, and determine if any of the metabolic enzymes produced by the pathogen might be potential antibacterial targets. This aim was accomplished through genome scale metabolic modeling of *B. pseudomallei* under different external conditions, including all nutrients that could be consumed by the model and only the nutrients available in culture media. Using this approach, candidate chokepoints were identified, then knocked out the genes encoding chokepoint enzymes *in silico* under the different nutrient conditions, and examined the effect of each knockout on the metabolic network. The result of this analysis was five candidate antibacterial targets, demonstrating the utility of genome-scale metabolic modeling methods for *in silico* studies of pathogen metabolism and for drug target identification in *B. pseudomallei*.

## 2. Materials and Methods

### 2.1 Metabolic pathway reconstructions and annotation curation

Pathway genome databases (PGDBs) for *B. pseudomallei* strains MSHR668 and K96243 were obtained through the Pathway Tools software (version 18.5) from the PGDB registry [35]. We found that the original annotation of *B. pseudomallei* K96243 identified many fewer coding sequences than that of MSHR668, so we re-annotated the original complete genome sequences of both strains using the RAST system [36] to better compare them [37]. The RAST-annotated genome sequences were loaded into Pathway Tools, using the PathoLogic component to predict the metabolic pathways [38]. For each genome, the set of protein coding sequences from the original annotation was compared to those from the RAST annotation, using blast to identify coding sequences in common between the two annotations, and to identify coding sequences that were missing from each annotation. For the MSHR668 genome, RAST annotation identified 247 fewer coding sequences than the original annotation. However, there were some predicted coding sequences with annotated functions in the RAST annotation that were not present in the original, so these were added to the original PGDB for MSHR668. The RAST annotation of the K96243 genome identified 1,317 more protein coding sequences than the original annotation, so the PGDB created from the RAST annotation was used as the starting point. Coding sequences from the original annotation that were not present in the RAST annotation of K96243 were added to the PGDB that was created from the RAST annotated genome.

### 2.2 Chokepoint Identification

Chokepoints were identified in each PGDB using the chokepoint reaction finder in Pathway Tools. All reactions were included except those found in human.

### 2.3 Flux balance analysis

For each metabolic network reconstruction, flux balance analysis (FBA) was performed using the MetaFlux module within the Pathway Tools [39]. Development FBA models were constructed iteratively to determine the compounds that each model could use and produce. This was accomplished by trying all compounds in the PGDB as biomass metabolites, nutrients and secretions in the various compartments (cytosol, periplasmic space and extracellular). Each model was refined iteratively, first identifying the specific biomass components that could be produced, then trying all compounds as nutrients and secretions, then specifying the biomass metabolites and nutrients and trying all compounds as secretions.

Once the nutrients, secretions and biomass components that could be consumed or produced by the metabolic networks were determined, the log file produced by MetaFlux was examined and problematic reactions were fixed, if possible. Most of the problematic reactions were unbalanced due to missing chemical formulas of one or more metabolites. A few of these reactions were corrected by copying the missing structures from other PGDBs. For example, the structures of D-ribose, D-glucuronate, D-glucose and some other compounds were copied from the more highly curated *Escherichia coli* K12 substr. MG1655 PGDB. Many of the reactions with compounds that were lacking chemical formulas were generic so no suitable chemical structure could be found or created. Reactions involving starch, glycogen, and glucans with variable lengths and non-numeric stoichiometries, could not be balanced. Other reactions were missing H+ or H2O on one side or the other, and the addition of the missing compound balanced the reaction. However, there were a small number of reactions that could not be fixed and these were marked as unbalanced. Once all reactions that could be fixed were corrected in the PGDB, MetaFlux was run again in development mode to identify additional biomass metabolites, nutrients, and secretions. Once the set of biomass metabolites was constant, the nutrients that could be consumed by the model were determined, followed by identification of the secretions produced by the model. The result of this process was a final unconstrained FBA model.

To mimic the nutrient conditions in culture, only the estimated set of ingredients present in LB medium were included as nutrients in the FBA model. LB medium includes as its main ingredients tryptone [40] and yeast extract [41]. Tryptone provides peptides and peptones, which are good sources of amino acids (http://www.sigmaaldrich.com/analytical-chromatography/microbiology/basic-ingredients/protein-sources.html); yeast extract provides vitamins, nitrogenous compounds, carbon, sulfur, trace elements and minerals [42]. The LB media composition reported in a *Bacillus subtilis* modeling study [43] was used as a starting point for this study. Starting with the nutrient set that mimicked LB media [43], all compounds were included in the try-biomass set for the cytosol and periplasmic cellular compartments to see which biomass metabolites could be produced given only the nutrients present in LB media. Once a stable set of nutrients was determined, iterations were performed to determine the stable set of biomass metabolites that could be produced. Then all compounds were tried as secretions in the cytosol and periplasmic compartments to determine the compounds that could be secreted by the model.

A list of nutrients that might be available to *B. pseudomallei* while residing inside host cells during infection was compiled by searching the literature for infection studies involving *B. pseudomallei* and host cells. Gene expression studies of other intracellular pathogens and host cells during infection were also considered. A development FBA model mimicking infection conditions was constructed as described above for the LB media model.

### 2.4 Essential gene and candidate drug target identification

To reduce the set of potential drug targets, candidate chokepoint enzymes were compared against the list of essential genes and all of the drug targets in DrugBank. Essential genes in the MSHR668 genome were determined by blasting all protein coding sequences (amino acid format) against the Database of Essential Genes [44] using the blastp program with an E-value cut-off of 1e - 10 and 70% identity as thresholds. Candidate drug target sequences were determined by blasting all protein coding sequences in the MSHR668 genome (as both nucleotide and amino acid formats) against the complete set of DrugBank targets [45], using an E-value cutoff of 0.005 and a threshold identity of 70% to select likely targets.

### 2.5 In silico knockout experiments

Knockout experiments were performed *in silico* through the Pathway Tools MetaFlux module [39]; the chokepoint genes were knocked out one at a time and the effects on total biomass flux through the metabolic network were noted.

### 2.6 Network visualization

For each PGDB, an.sbml file was exported from the Pathway Tools and loaded into Cytoscape [46] version 3.1.1 for visualization and comparison of network features.

## 3. Results

### 3.1 General features of B. pseudomallei genomes and metabolic networks

The general characteristics of each PGDB and metabolic network for B. pseudomallei MSHR668 and K96243 are listed in Table 1.

**Table 1.** Features of *B. pseudomallei* PGDBs and metabolic networks

| Genome<br>(Pathway Tools PGDB) | Curated MSHR668<br>(original + RAST) | Curated K96243<br>(RAST + original) |
| --- | --- | --- |
| Coding sequences | 7133 | 7045 |
| Pathways | 339 | 387 |
| Enzymatic reactions | 1870 | 1995 |
| Transport reactions | 144 | 82 |
| Enzymes | 1666 | 1685 |
| Transporters | 292 | 38 |
| Compounds | 1397 | 1578 |
| Metabolic Network |  |  |
| Nodes (metabolites) | 4295 | 4219 |
| Edges (reaction steps) | 10860 | 9747 |
| Chokepoints |  |  |
| Producing | 444 | 506 |
| Consuming | 419 | 498 |
| Candidate | 479 | 348 |
| Dead-end reactants | 166 | 217 |
| Dead-end products | 72 | 118 |

The *B. pseudomallei* MSHR668 and K96243 genomes contained similar numbers of coding sequences, pathways, enzymatic reactions and enzymes. Differences between the PGDBs were noted in the numbers of transporters, transport reactions and compounds. The K96243 PGDB contained fewer transporters and transport reactions and more compounds than MSHR668. In terms of the metabolic network characteristics, both networks contained similar numbers of nodes (representing metabolites), while the MSHR668 network had more edges (reaction steps) than K96243. This is likely due to the more extensive curation of the MSHR668 network that was performed during refinement of the metabolic network models (see Methods).

In a metabolic network, chokepoints are reactions that either uniquely produce or uniquely consume a metabolite. Inhibiting an enzyme that consumes a unique substrate may cause that metabolite to accumulate, and it may be toxic to the cell; inhibiting an enzyme that produces a unique product may starve the cell of an essential metabolite [31]. Identifying chokepoint enzymes in pathogens is a promising *in silico* approach to recognize potential metabolic drug targets. For example, analysis of *Plasmodium falciparum* metabolism revealed that 87.5% of proposed drug targets supported by evidence are chokepoint reactions [31]. However, to be a valid chokepoint, the metabolite in question must be balanced by a producing or consuming reaction and not be a deadend metabolite [31]. Table 1 compares the numbers of chokepoint reactions and dead-end metabolites that were identified in each PGDB. Overall the numbers were similar between the two PGDBs, and the lower numbers of dead-ends in the MSHR668 database were likely due to the more extensive curation performed on the MSHR668 PGDB (see Materials and Methods).

### 3.2 Flux balance analysis (FBA)

#### 3.2.1. Unconstrained FBA model

Given a set of nutrients for consumption, along with secretions and metabolites that can be produced, a FBA model predicts the steady-state flux rates of the metabolic reactions in an organism, and provides an estimate of the overall biomass flux. FBA was conducted on each metabolic network as described in the Materials and Methods section. It took several cycles of refinement to solve an initial unconstrained MSHR668 model. Even after several iterations, the K96243 unconstrained model did not reach a stable solution, likely because there were missing transporter-encoding genes (and possibly other genes) in the PGDB, indicating that the annotation needed more curation. Since the unconstrained MSHR668 model reached a stable solution after performing the initial network curation steps suggested by MetaFlux, only this model was analyzed further. The initial unconstrained FBA model of the MSHR668 metabolic network (Table 2) included all possible biomass compounds that could be produced, all nutrients that could be consumed, and had no weights imposed on the biomass metabolites and no constraints imposed on the nutrients. [Supplementary materials, S1_Final_unconstrained_model_inputs.pdf, S2_Final_unconstrained_model_solution.pdf].

#### 3.2.2. LB media FBA model

To mimic the conditions that *B. pseudomallei* experiences in culture, only the nutrients present in LB media [43] plus glycerol were included as inputs to the LB media FBA model (Table 2). In this model, constraints were included on some of the nutrients (ADP, Pi, proton and glycerol). [Supplementary materials, S3_Final_LB_model_inputs.pdf and S4_Final_LB_model_solution.pdf].

**Table 2:** Characteristics of the MSHR668 unconstrained and LB media models

| Model | Total rxns | Rxns carrying flux | Biomass metabolites produced | Nutrients consumed | Secretions | Biomass flux |
| --- | --- | --- | --- | --- | --- | --- |
| Unconstrained | 3193 | 1619 | 1403 | 667 | 10 | 15000.00 |
| LB media | 3213 | 1025 | 282 | 47 | 0 | 0.079412 |

#### 3.2.3. Host cell infection model

In addition to the unconstrained and LB media models, the original plan for this study included the development of a model of *B. pseudomallei* metabolism that mimics infection conditions. However, there was very little information available on the growth requirements of *B. pseudomallei* inside human macrophages. In addition, no comprehensive studies have been performed to identify the complete list of host cell nutrients that are available to *B. pseudomallei* during infection. Most studies of the nutritional requirements of intracellular pathogens growing inside host cells have been performed on *Legionella pneumophila* [47,48], which can grow and replicate similarly in human macrophages and amoebae [49]. Growing in both human macrophages and amoebae, *L. pneumophila* utilizes amino acids as its main sources of carbon, nitrogen and energy; *L. pneumophila* obtains amino acids from the host through proteasomal degradation [48]. However, glucose is also used to feed central metabolism under both culture and infection conditions [50].

Comparing the nutrients provided by the LB media model to the nutrients used by *L. pneumophila* in culture and during infection of amoebae [47], the only difference was the carbohydrate carbon source: glycerol *(B. pseudomallei* LB media [51]) vs. glucose *(L. pneumophila* AYE media [50] and in amoebae [47,50]). When glycerol was replaced by glucose in the LB media model of *B. pseudomallei,* no FBA solution was found [data not shown]. Several possible explanations for this result are presented in the Discussion.

While the specific carbon requirements of *B. pseudomallei* in either human macrophages or amoebae have not been determined, one study produced whole-genome tiling array expression data to assess *B. pseudomallei* transcriptional responses under 82 different conditions, including infection [52]. From their supplemental table S2, a list of metabolic genes expressed in the infection conditions was used to infer the potential nutrients consumed by *B. pseudomallei* during infection. Additional candidate host cell nutrients were identified from the literature, focusing on studies of intracellular pathogen-mammalian host infections. The nutrients identified as potential carbon sources for intracellular survival of various pathogens included aromatic compounds, such as benzoate and phenylacetic acid and related derivatives [53], sugar acids like gluconate, galactonate, glucuronate, and galacturonate [54], ribo- and deoxyribonucleosides, hexuronates [55], glutathione [56], glucose 6-phosphate [57,58], glycerol-3-phosphate [59]. The complete list of potential host cell nutrients is in the [Supplementary material, S5_Nutrients_infection_model.pdf] file.

FBA was performed for a *B. pseudomallei* MSHR668 model that included the list of candidate host cell nutrients identified as above. However, some of the nutrients were not present in any reactions in the MSHR668 PGDB. When the rest of the compounds were included as nutrients in addition to glycerol, none of them were consumed by the model, but glycerol was consumed and biomass was produced. When glycerol was excluded from the nutrient list, none of the other nutrients were consumed and the model was not solvable [data not shown].

### 3.3 Metabolic chokepoints and candidate drug targets

To narrow down the list of metabolic chokepoints, which represented candidate drug targets, essential genes and genes with sequence homology to existing DrugBank targets were identified in the MSHR668 genome. This analysis identified 34 chokepoint genes that were also essential genes and DrugBank targets (Figure 1, Table 3).

**Figure 1.**
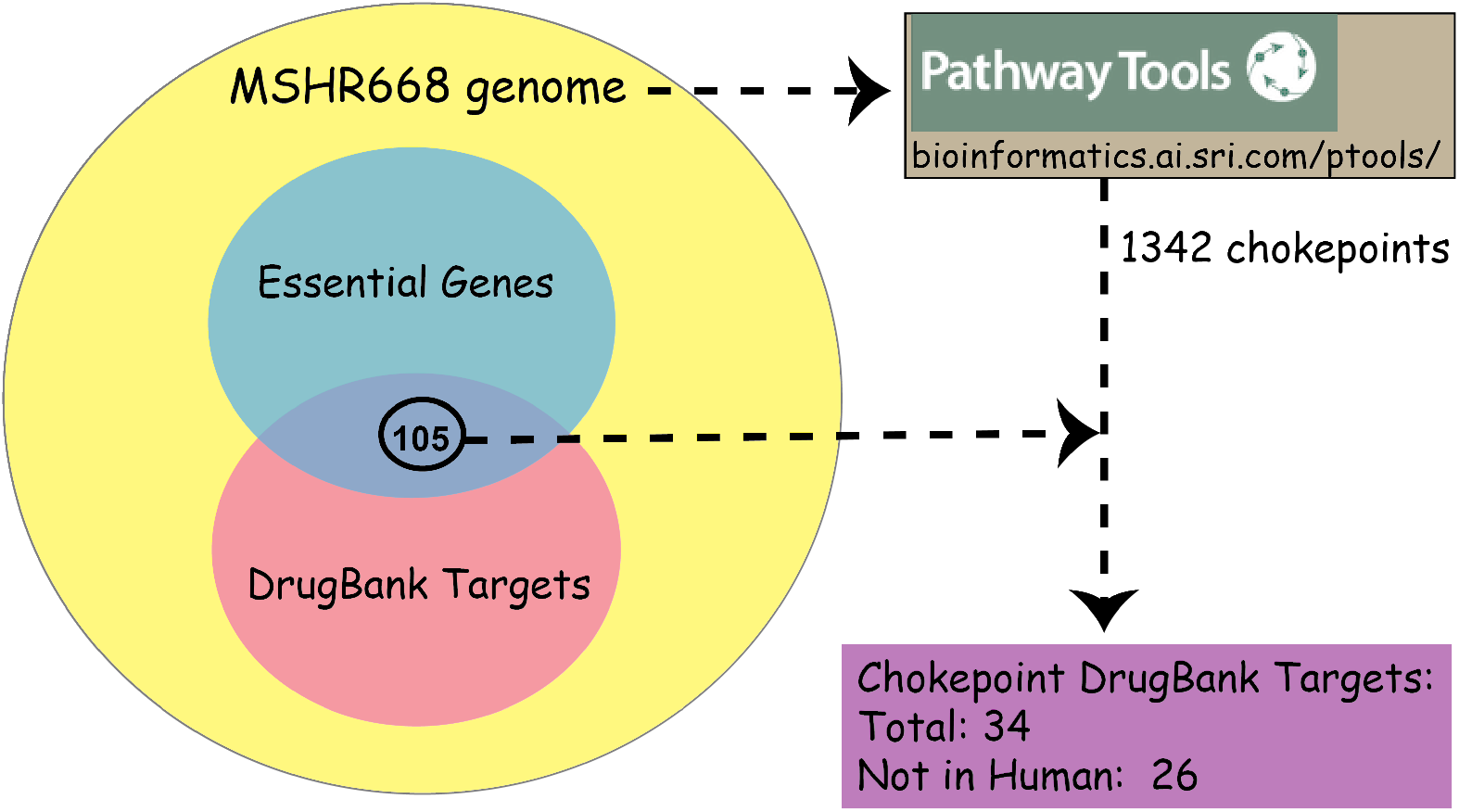
Process for identifying candidate metabolic enzyme drug targets (chokepoints) in the *B. pseudomallei* MSHR668 genome.

**Table 3.**
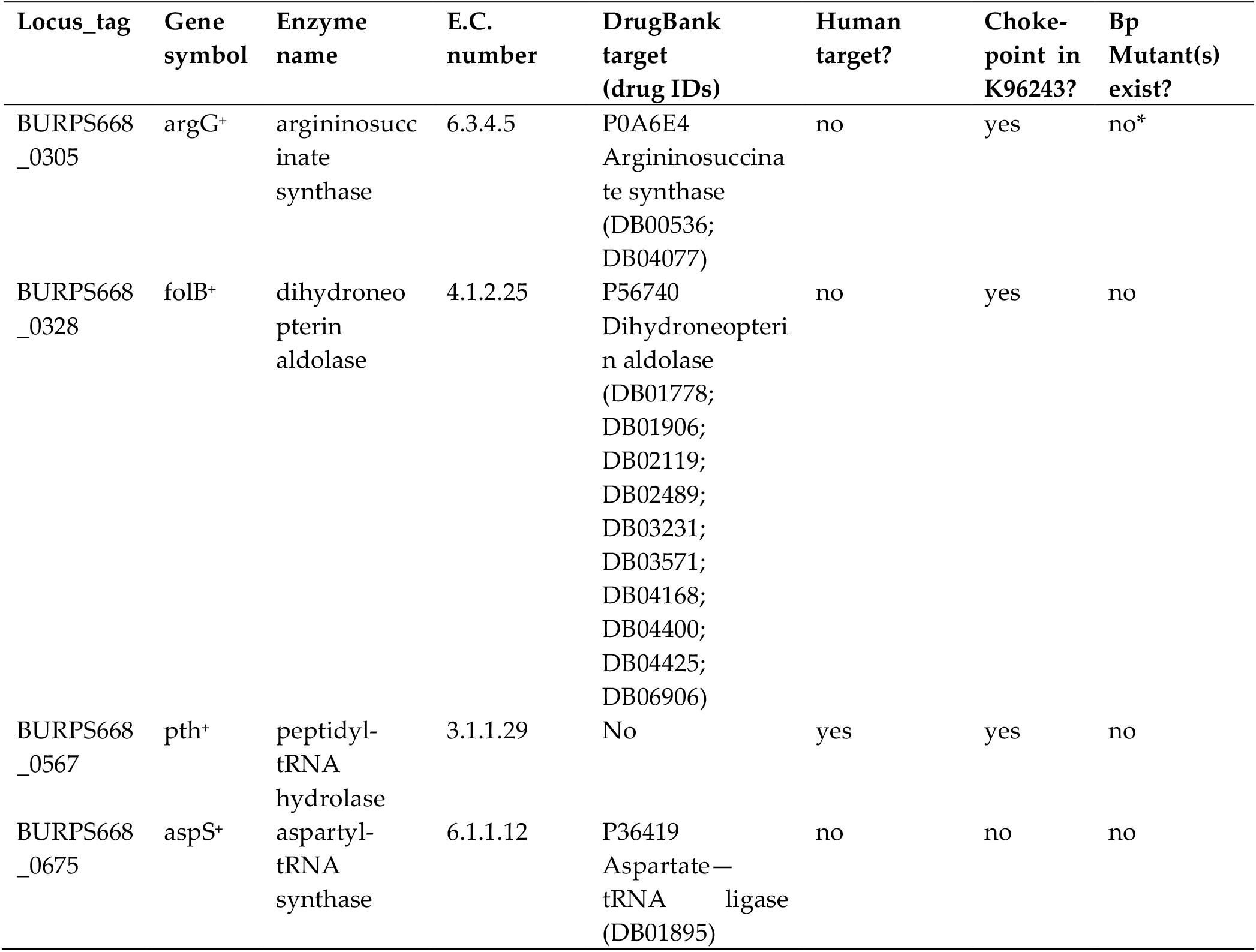
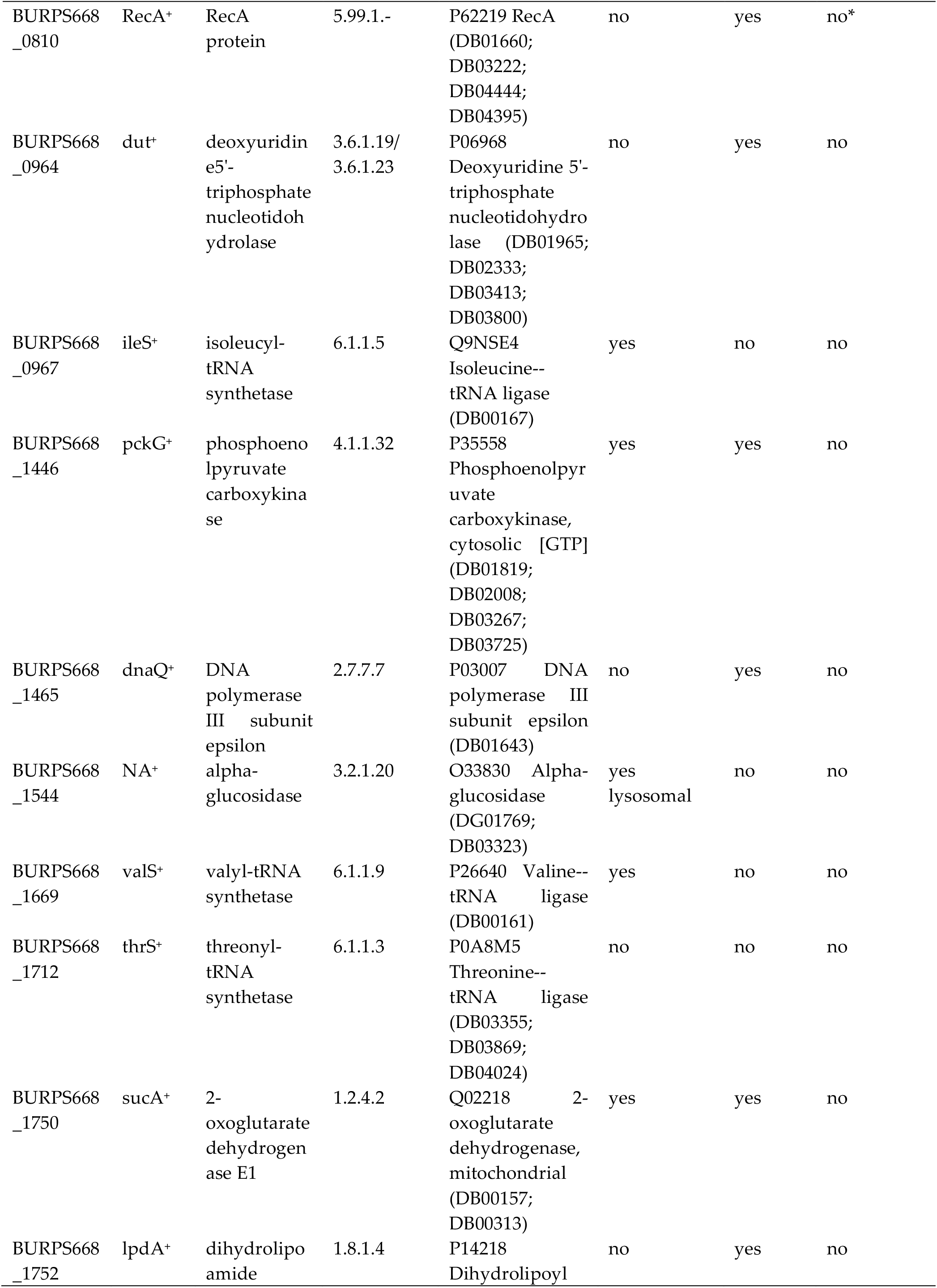
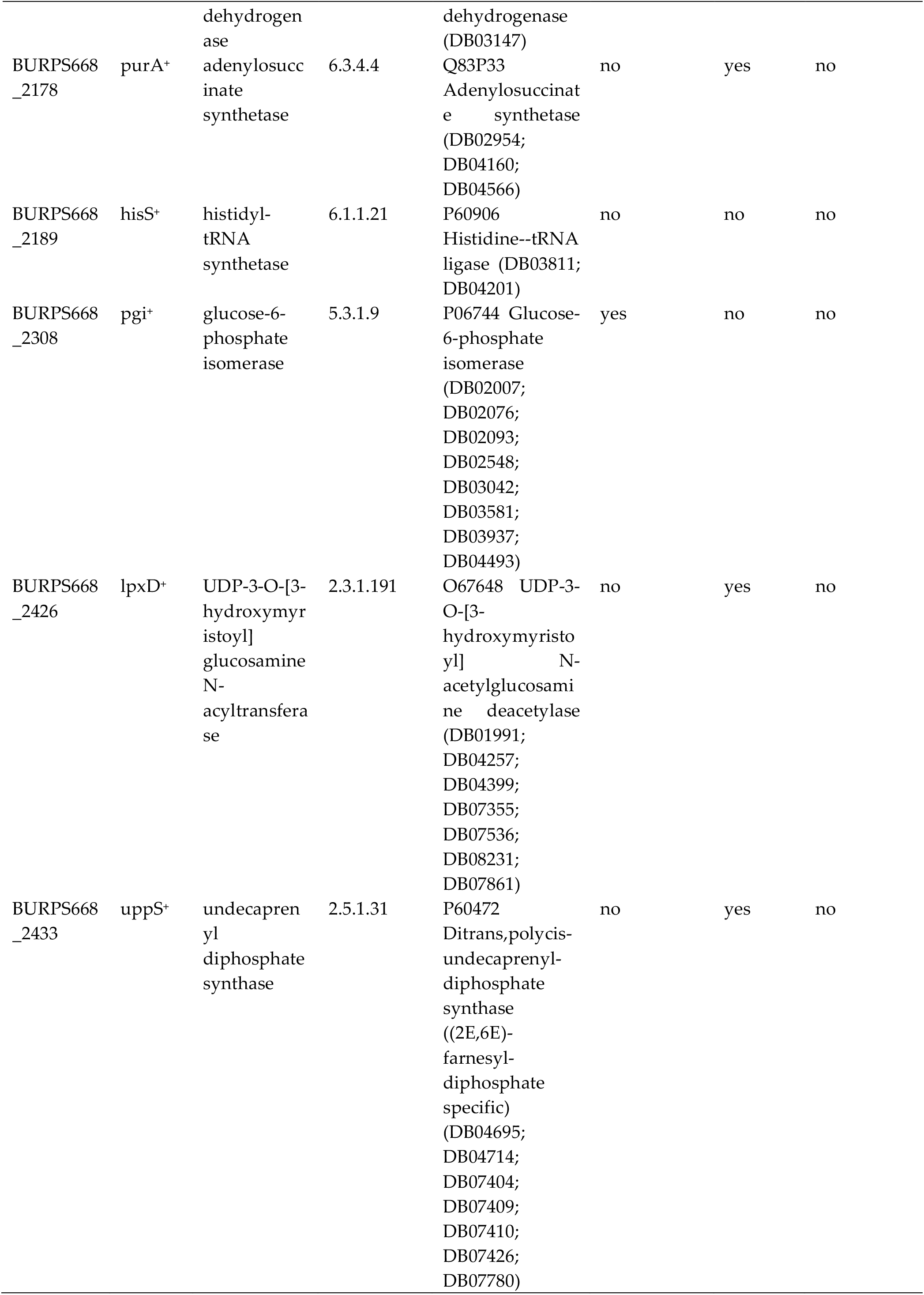
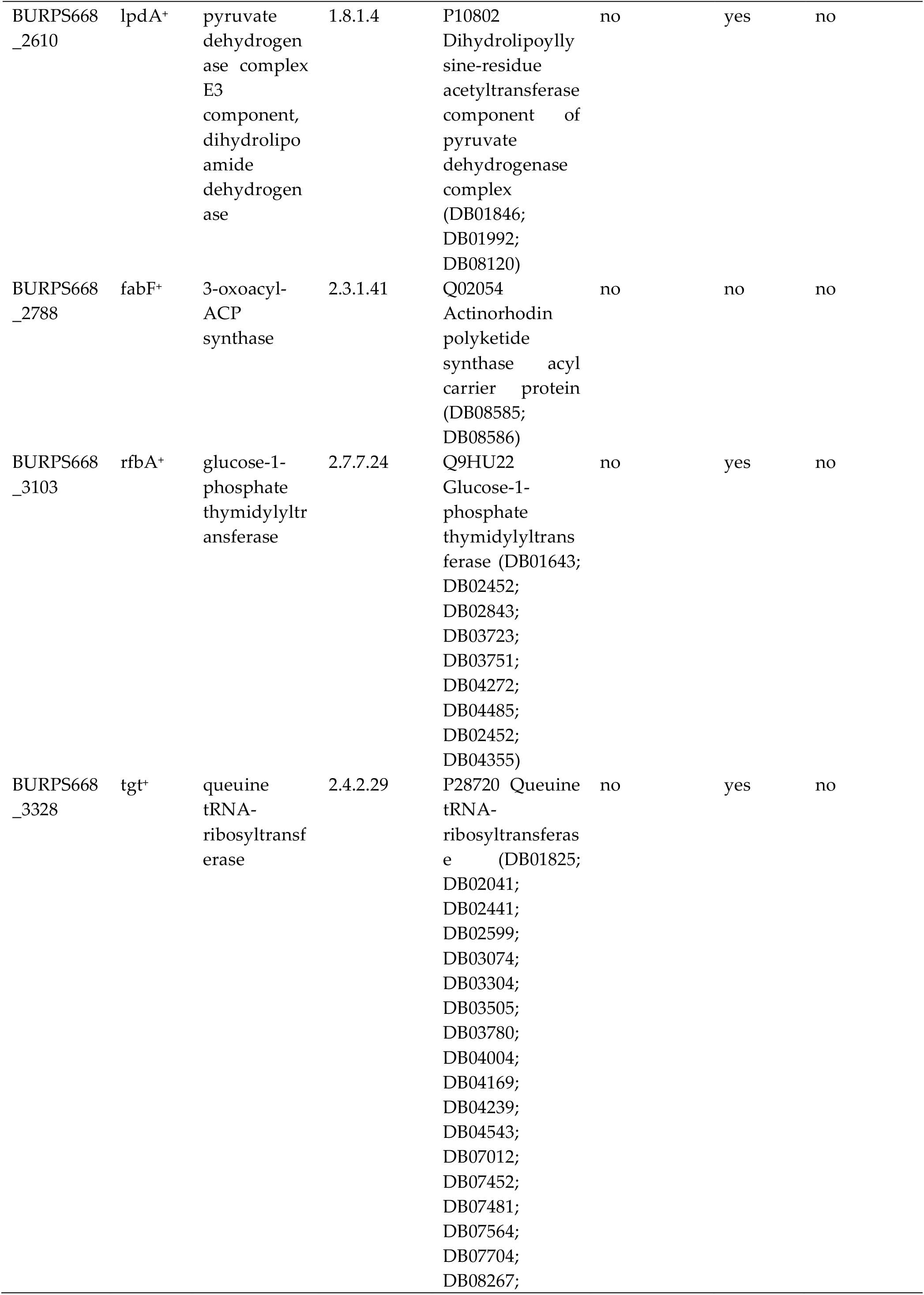
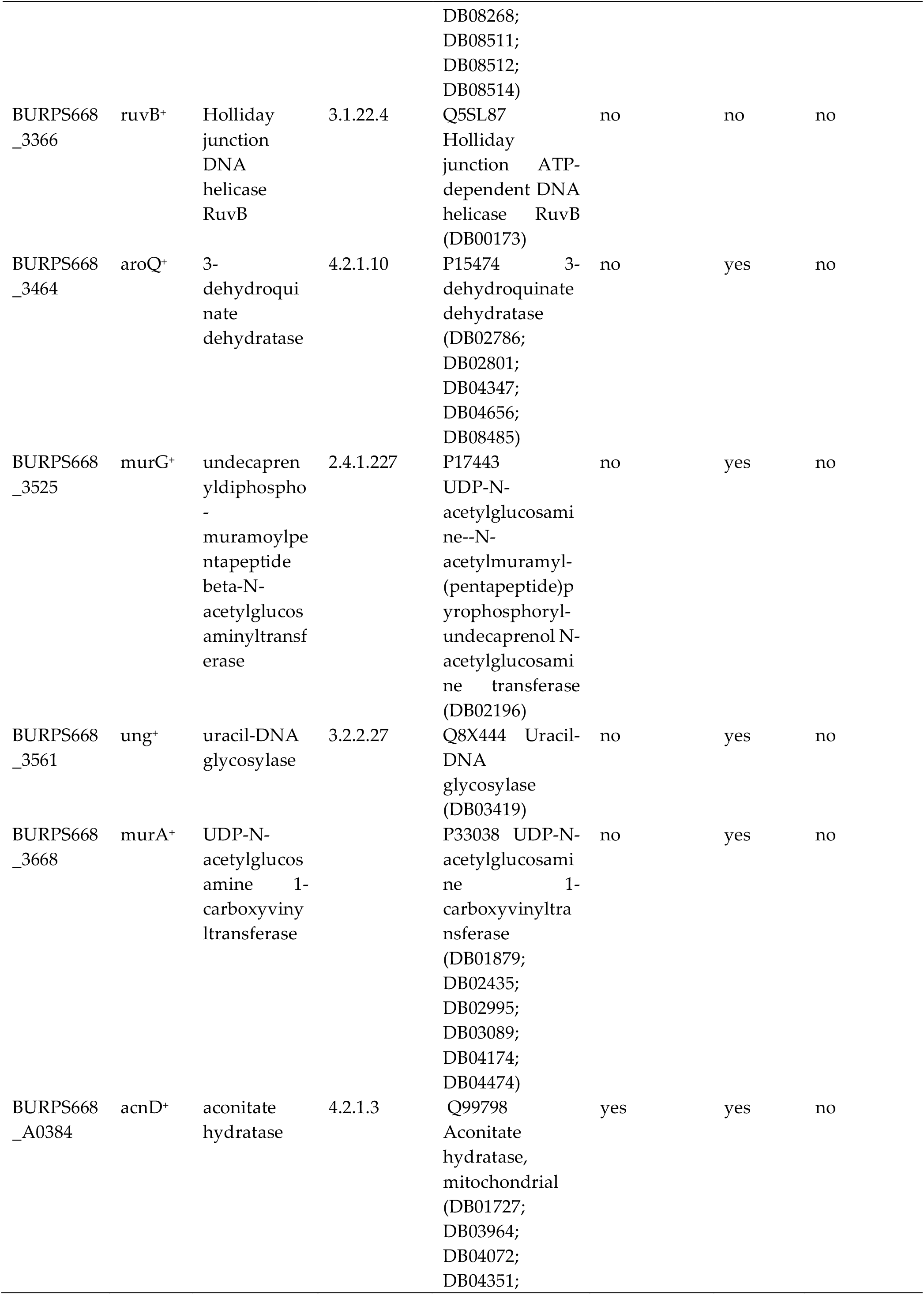
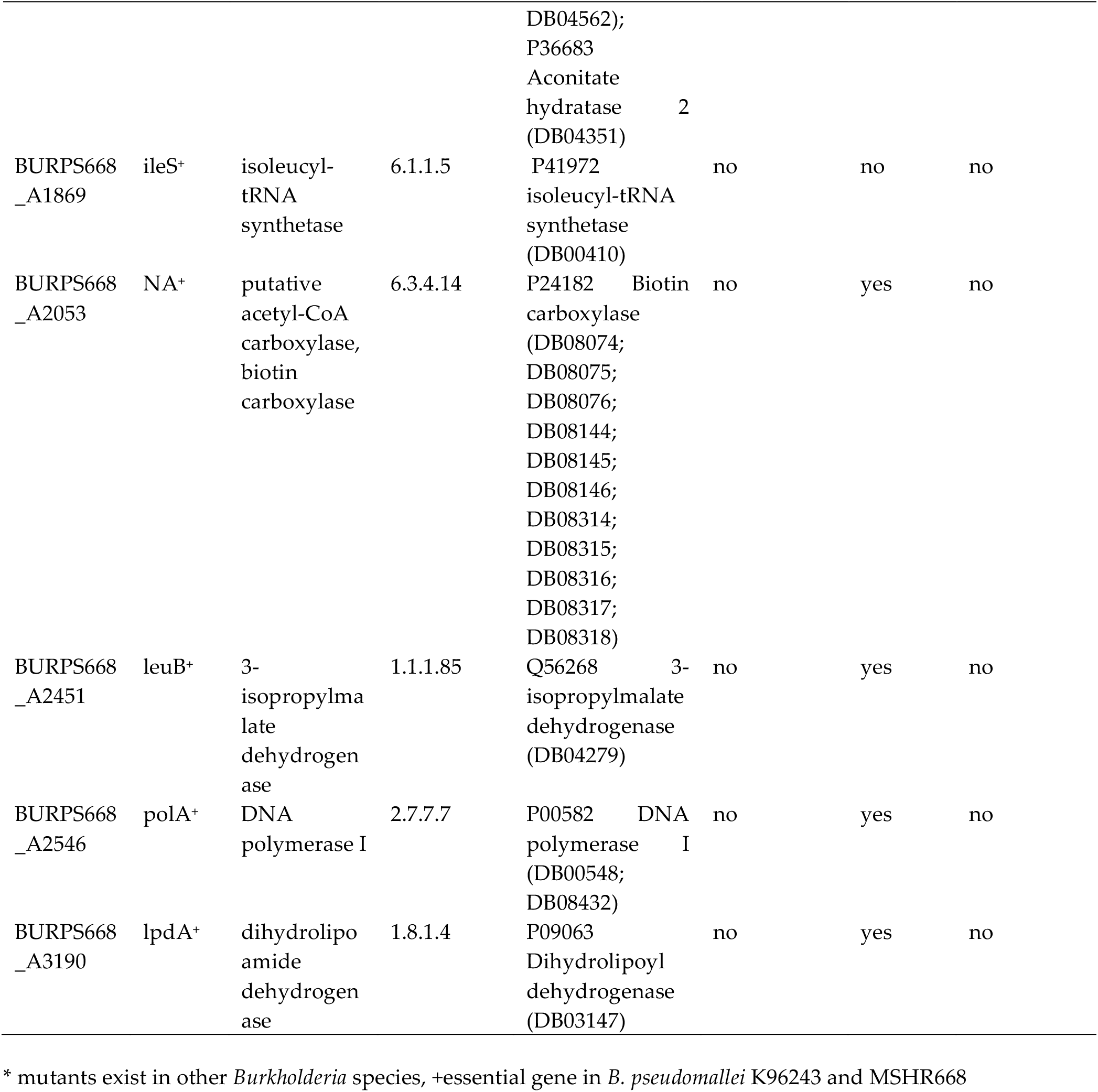
Chokepoint genes that encode candidate metabolic enzyme drug targets in *B. pseudomallei* MSHR668

Eight of the targets in Table 3 were also DrugBank targets in human; DrugBank was searched for the remaining twenty-six targets, showing that they also occur in other bacteria.

*In silico* knockout experiments were performed with the MetaFlux module of Pathway Tools to test the effect of inhibiting each chokepoint enzyme on *B. pseudomallei* growth in both the unconstrained and LB media models. Results (Table 4) show that knockout of BURPS668_3328 (tgt) and BURPS668_A2451 (*leuB*), eliminated the biomass flux in the unconstrained model, while knockout of BURPS668_2426 *(lpxD),* BURPS668_2433 (uppS), and BURPS668_3525 *(murG)* eliminated the biomass flux in the LB media model. Knockout of BURPS668_2433 *(uppS),* and BURPS668_3525 (*murG*) decreased the biomass flux in the unconstrained model, but did not eliminate it.

**Table 4.**
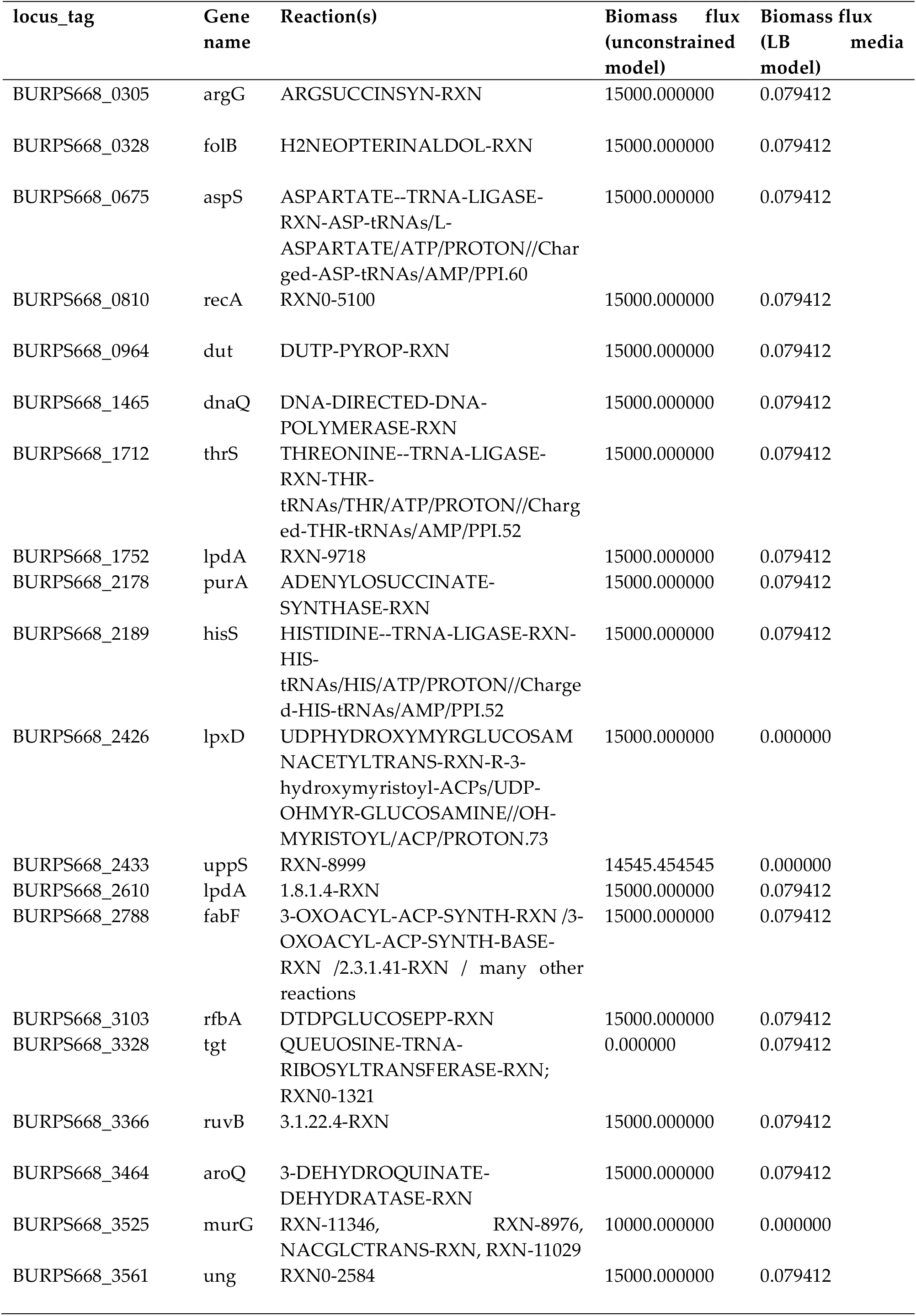
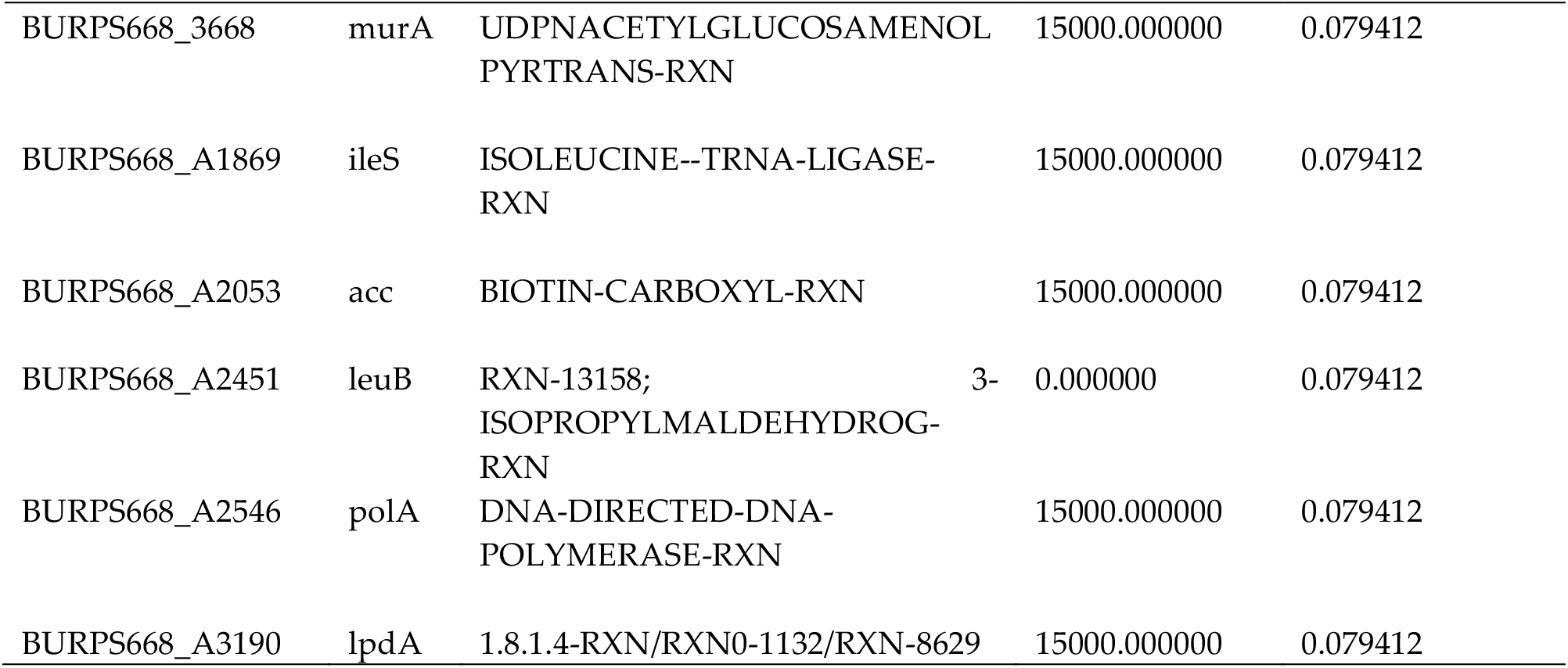
Results of *in silico* knockout experiments on twenty-six chokepoint reactions in *B. pseudomallei*

The overall biomass fluxes were different in the two models. The unconstrained model had a much greater total biomass flux than the LB media model. This is likely because the unconstrained model included more nutrient inputs than the LB media model.

## 4. Discussion

### 4.1 Links between metabolism and virulence

In order to colonize the host, establish an infection, and proliferate, pathogens employ various strategies, often involving links between metabolic pathways and virulence genes. Although current knowledge regarding the connections between metabolism and virulence is limited [60], this topic is becoming an increasing focus for host-pathogen studies. Some general links between metabolism and virulence include regulatory connections between specific metabolites and virulence gene expression [61–65], metabolic requirements for adaptation of the pathogen to the host niche [59,60,66,67], and carbon catabolite repression [68]. More detailed information on the topic of metabolism and virulence can be found in reviews [59,60,66,67,69–74].

Inside host cells, the survival of pathogenic bacteria depends on their acquisition of nutrients and carbon sources, such as carbohydrates, lipids, glycolipids, dicarboxylic acids and amino acids, from their host environment [59,60,66,67,71,75]. Preferred carbon sources vary among intracellular pathogens, and the types of nutrients available in the host cell cytosol may determine the cell-type specificities of different intracellular pathogens [76]. For example, many bacteria prefer hexoses, like glucose, as sources of carbon and energy. These sugars are catabolized through glycolysis, the pentose phosphate and Entner-Doudoroff pathways [75]. Some bacteria lack the glycolysis pathway and preferentially metabolize glucose via the Entner-Doudoroff pathway [77], while others lack both glycolysis and Entner-Doudoroff pathways and live on pyruvate that they obtain from the host cell cytosol [78].

### 4.2 Metabolic potential of B. pseudomallei MSHR668

We previously reported that *B. pseudomallei* MSHR668, K96243 and 1106a have abundant capabilities to metabolize hexoses, including the complete sets of genes encoding the glycolysis, pentose phosphate cycle, and Entner-Doudoroff pathways. They also have several pathways for metabolism of pyruvate to acetyl-CoA, acetate and ethanol [37]. In general, *B. pseudomallei* as a species has a very diverse set of metabolic capabilities, likely a reflection of its ability to live both in the natural environment and in hosts.

The metabolic power of *B. pseudomallei* has been targeted in mutation studies to address the roles of various metabolic genes in virulence. Mutation of various metabolic genes that affect cell growth results in attenuation of *B. pseudomallei* virulence; these genes include phosphoribosylformylglycinamidine cyclo-ligase *(purM)* [79], aspartate-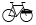semialdehyde dehydrogenase *(asd)* [80], acetolactate synthase *(ilvI)* [21], dehydroquinate synthase *(aroB)* [20], chorismate synthase *(aroC)* [17], phosphoserine aminotransferase *(serC)* [81], phosphoribosylglycinamide formyltransferase 1 *(purN)* and phosphoribosylformylglycinamide cyclo-ligase *(purM)* [18], two phospholipase C enzymes [82], disulfide oxidoreductase *(dsbA)* [83]. Targeted mutation of *purM,* which encodes aminoimidazole ribotide, a precursor of de novo adenine and thiamine biosynthesis, predictably causes a deficiency in adenine and thiamine biosynthesis [79]. Aspartate-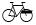semialdehyde dehydrogenase *(asd)* mutants cannot synthesize diaminopimelate for cell wall biosynthesis [80], while acetolactate synthase *(ilvI)* mutants cannot synthesize the branched chain amino acids isoleucine, valine, and leucine [21]. Dehydroquinate synthase *(aroB)* mutants are defective in the shikimate pathway for chorismate biosynthesis, and addition of the aromatic compounds tyrosine, tryptophan, phenylalanine, PABA, and 2,3-dihydroxybenzoate is required to restore the growth in minimal medium [20]. Chorismate synthase (*aroC)* mutants are also defective in aromatic compound synthesis and cannot grow without the addition of aromatic compounds to the media [17]. Phosphoserine aminotransferase *(serC)* mutants are defective in serine and pyridoxal 5-phosphate biosynthesis and require minimal medium supplemented with serine for normal growth [81].

### 4.3 Vaccine studies

Using live attenuated bacteria as vaccines can be effective in preventing disease, as attenuated bacteria may still be able to replicate in the host and may contain immune-stimulatory epitopes that are not found in subunit or heat-inactivated vaccines [84]. Many of the *B. pseudomallei* mutants described above have already been tried as attenuated vaccines with mixed results. Vaccination with the attenuated *asd* mutant protected BALB/c mice against acute melioidosis, but did not protect against chronic melioidosis [80]. Vaccination with the attenuated *ilvI* mutant of *B. pseudomallei* protected BALB/c mice against a challenge with a virulent strain [21]. In mice vaccinated with the *aroB* mutant, the time to death following challenge with the virulent K96243 strain was a bit longer than in unvaccinated mice, but all mice eventually died [20]. The *aroC* mutant was unable to persist in vaccinated BALB/c mice long enough to elicit protective immunity, however C57BL/6 mice were protected against challenge with a virulent strain [17]. Intraperitoneal vaccination of BALB/c mice with a *serC* mutant resulted in higher levels of survival after challenge with K96243 virulent strain [81]. While immunization of mice with attenuated *B. pseudomallei* mutants has resulted in the induction of protective immunity in some cases, sterile immunity was rarely reported (reviewed by [85]). Also, the live attenuated vaccine model may not be the best solution for the prevention of melioidosis, because an attenuated mutant might revert to virulence, and might also establish a latent infection [85].

### 4.4 Identification of antimicrobial therapeutics

An alternative avenue to combat melioidosis is through the development of novel antimicrobial therapeutics. Regardless of the specific metabolic capabilities possessed by a pathogen, essential nutrient acquisition and utilization mechanisms are proving to be good potential therapeutic targets, as inhibition of these targets might deprive the pathogen of needed substrates for growth and replication inside host cells [31]. There are currently 699 *B. pseudomallei* genomes available at the National Center for Biotechnology Information (NCBI, http://www.ncbi.nlm.nih.gov/genome/476). While some of these entries represent re-annotations of previous submissions, and some genomes represent colony morphology variants of the same strain, an impressive number of individual genomes are available to use with computational approaches to identify new therapeutic targets. With this large number of available genomes, a core genome approach [86,87] could be used to identify potential metabolic enzyme targets that are present in all sequenced *B. pseudomallei* genomes.

*In silico* methods for the identification of therapeutic targets in bacterial pathogens include comparative genomics-based approaches, such as identification of essential genes specific to the pathogen, and techniques based on metabolic pathway analysis and metabolic network modeling. The more robust approaches use a combination of comparative genomics and metabolic pathway analysis. These approaches have been used to identify essential gene targets in *Mycoplasma genitalium* [88] and *Mycobacterium ulcerans* [89]. Another method, subtractive target identification, involves identification of enzymes in the metabolic pathways of the pathogen, and comparing them to human proteins to identify pathogen enzymes that are not found in human. A list of likely targets is compiled by focusing on enzymes in pathways that are usually essential for pathogen growth and survival, like lipid metabolism, carbohydrate metabolism, amino acid metabolism, energy metabolism, vitamin and cofactor biosynthetic pathways and nucleotide metabolism. This approach has been used to identify putative targets in *M. tuberculosis* [90], MRSA [91–93], as well as a collection of other bacterial pathogens [94].

Methods for therapeutic target discovery based on metabolic pathway analysis and metabolic network modeling have become very popular in the last ten years. Numerous studies have identified candidate drug targets in various bacterial [27,34,95–106], fungal [106] and protosoan [31,106–109] pathogens using a variety of methods to analyze metabolic pathways and networks. One method employs chokepoint analysis to identify metabolic enzymes that are critical to the pathogen, because they uniquely consume and/or produce certain metabolites. Chokepoint analysis has been used to identify candidate metabolic enzyme targets in various pathogen genomes [27,31,34,99,107,108,110,111]. However, no studies to date have used this approach to identify potential drug targets in *B. pseudomallei* metabolic networks.

#### 4.4.1 Identification of metabolic chokepoints

For this study, metabolic chokepoints were identified in the curated *B. pseudomallei* MSHR668 metabolic network using the Pathway Tools software [35]. Table 3 lists the chokepoint enzymes identified in the metabolic networks of *B. pseudomallei* MSHR668 and K96243. Twenty-four of the *B. pseudomallei* chokepoints were not indicated as human targets in DrugBank, and therefore represented good candidate therapeutic targets against melioidosis. Six of the chokepoints in Table 3 were aminoacyl-tRNA synthetases, which are likely good targets as they are critical enzymes involved in protein translation. These chokepoints included aspartyl-, threonyl-, histidyl-, valyl- and isoleucyl-tRNA synthetase (2 copies). Aspartyl-tRNA synthase *(aspS)* is an essential gene target in *M. tuberculosis* [112]. Threonyl-tRNA synthetase *(thrS)* inhibitors have been identified [113] and shown to have anti-malarial activity against *Plasmodium falciparum* [114]. Isoleucyl-tRNA synthetase *(ileS)* is a well-documented bacterial target [115–117]. The antimicrobial drug, mupirocin (pseudomonic acid), selectively inhibits bacterial isoleucyl-tRNA synthetase without inhibiting its human homolog [114]. However, resistance is seen in bacteria that possess an isoleucyl-tRNA synthetase that is similar to eukaryotic versions [118]. Histidyl-tRNA synthetase *(hisS)* has been explored as a target in *Trypanosoma cruzi* [119]. The chokepoint enzyme queuine tRNA-ribosyltransferase (tgt) incorporates the wobble base queuine into tRNA, and is also a target in *Zymomonas mobilis* and *Shigella* [120,121]

Several *B. pseudomallei* chokepoint enzymes are likely involved in DNA-related processes. Two of these enzymes, encoded by the *recA* and *dnaQ* genes, are involved in the SOS pathway [122], which mediates the bacterial response to DNA damage. Activation of the SOS response by ciprofloxacin induces mutations, which can lead to fluoriquinolone resistance [123]. The RecA protein is a target for antibacterial drug discovery in *M. tuberculosis* [124] and *Mycoplasma hyopneumoniae* [96], and has been proposed as a specific target for reducing the evolution of antimicrobial resistance [125]. The chokepoint enzyme deoxyuridine 5’-triphosphate nucleotidohydrolase (dUTPase) prevents incorporation of uracil into DNA and is important for DNA integrity [126]. dUTPase is a potential antimalarial drug target against *P. falciparum* [127,128]. Holliday junction DNA helicase *(ruvB)* participates in homologous recombination and repair of replication forks, and is therefore essential for bacterial growth. Holliday junction processing components were previously identified as targets for antimicrobials in *E. coli* [129] and *Neisseria gonorrhoeae* [130]. The chokepoint enzyme uracil-DNA glycosylase *(ung)* has a role in uracil excision repair, and is a candidate anti-malarial drug target [131], as well as a potential target to control growth of GC-rich bacteria such as *Pseudomonas aeruginosa* and *Mycobacterium smegmatis* [132]. DNA polymerase I was identified as a chokepoint in *B. pseudomallei.* Putative inhibitors of DNA polymerase I (*polA*), and subsequently DNA synthesis, have been explored as possible antimicrobials [133,134].

Some chokepoint enzymes in *B. pseudomallei* have annotated functions in the biosynthesis of cell wall components. One of these, UDP-3-O-[3-hydroxymyristoyl] glucosamine N-acyltransferase *(lpxD),* catalyzes the third step in the lipid A biosynthesis pathway [135]. The lipid A component of bacterial LPS is of particular interest because it is essential for cell viability and is highly conserved [136]. This pathway is a target for new antibacterial therapeutics in *Escherichia coli* [137]. Another chokepoint involved in cell wall synthesis is undecaprenyl diphosphate synthase *(uppS),* which catalyzes the synthesis of a polyisoprenoid essential for both peptidoglycan and cell wall teichoic acid synthesis. UppS is a critical enzyme required for bacterial survival, and is an antibacterial target in *Staphylococcus aureus* [138], *Bacteroides fragilis, Vibrio vulnificus, E. coli* [139] and *H. pylori* [140]. Several classes of compounds that inhibit UppS function have been discovered [138,141,142]. Two additional *B. pseudomallei* chokepoints, undecaprenyldiphospho-muramoylpentapeptide beta-N-acetylglucosaminyltransferase *(murG),* and UDP-N-acetylglucosamine 1-carboxyvinyltransferase (*murA*) are likely involved in peptidoglycan biosynthesis. MurG is the target of the antibiotic ramoplanin in *Staphylococcus aureus* [143]. Other potential inhibitors of MurG have been identified by high throughput screening [144]. A small molecule inhibitor of MurG that augments the activity of β-lactams against methicillin-resistant *Staphylococcus aureus* was recently identified [145]. MurA has been a popular target for the design of novel antibiotics, and several inhibitors of MurA have been identified that are active against various bacterial species [91,146–151]. The chokepoint enzyme glucose-1-phosphate thymidylyltransferase *(rfbA/rmlA),* involved in O antigen biosynthesis, is also a target in *Streptococcus pneumoniae* [152] and *Pseudomonas aeruginosa* [153].

The rest of the chokepoint enzymes in Table 3 are components of various biosynthesis pathways. These enzymes include argininosuccinate synthase *(argG),* which catalyzes the second to last step in L-arginine biosynthesis, and is associated with pathogenesis in the parasite *Leishmania donovani* [154], *Streptococcus pneumoniae* [155], and *B. cenocepacia* [156]. The chokepoint enzyme 3-isopropylmalate dehydrogenase *(leuB)* is the third enzyme specific to leucine biosynthesis in microorganisms [157], and has been investigated as an antibacterial target in *M. tuberculosis* [158]. The *B. pseudomallei* chokepoint dihydroneopterin aldolase *(folB*) is part of the tetrahydrofolate biosynthesis process, and is essential for growth and biomass production in *Acinetobacter baylyi, Bacillus anthracis, Francisella tularensis, F. tularensis* subsp. novicida strain U112, *Mycobacterium tuberculosis, Helicobacter pylori, Pseudomonas aeruginosa, Salmonella enterica* serovar Typhi and *Yersinia pestis* [95]. Three of the chokepoint genes identified in *B. pseudomallei* encoded lipoamide dehydrogenase, a component of the pyruvate dehydrogenase complex, which converts pyruvate to acetyl-CoA as part of central metabolism [159]. Lipoamide dehydrogenase is also a target in *M. tuberculosis*, where deletion drastically impaired the pathogen’s ability to establish infection in the mouse [160]. The mycobacterial version has only 36% identity with the human homolog. Lipoamide dehydrogenase is a target of drugs against trypanosomal infections [161]. Adenylosuccinate synthetase *(purA)* is also a chokepoint enzyme and potential therapeutic target that is involved in purine salvage in *Leishmania donovani* [162]. Chokepoint enzyme 3-dehydroquinate dehydratase *(aroQ)* is a component of the shikimate pathway for chorismate biosynthesis and is a target of known inhibitors in *M. tuberculosis*, *Enterococcus faecalis* and *Streptomyces coelicolor* [163–167].

The *B. pseudomallei* chokepoint enzyme 3-oxoacyl-ACP synthase (*fabF*), involved in fatty acid synthesis, is already an antibacterial target in *E. coli*, and a specific inhibitor, cerulenin, has been identified [168,169]. Another fatty acid synthesis chokepoint enzyme in *B. pseudomallei* was a biotin-dependent acetyl-CoA carbosylase. Biotin dependent carboxylases comprise a large group of enzymes that participate in a variety of cellular processes, including fatty acid metabolism, amino acid metabolism, carbohydrate metabolism, polyketide biosynthesis, urea utilization, etc. (reviewed by [170]). Acetyl-CoA carboxylase is comprised of two enzymes, biotin carboxylase and carboxyltransferase, and catalyzes the first committed step in fatty acid synthesis [171]. Acetyl-CoA carboxylase is an antimicrobial target in *M. tuberculosis* [172], *E. coli* [173,174], other bacteria and most living organisms (reviewed by [175]). All of the chokepoints in Table 3 are essential genes in *B. pseudomallei* MSHR668 and K96243, as determined by blasting the chokepoint enzyme sequences against essential gene sequences in the Database of Essential Genes [176] and by comparing to the list of essential genes previously identified in K96243 [177].

To determine if *B. pseudomallei* deletion mutants were available for each of the chokepoints in Table 3, searches of the internet, PubMed, and the Burkholderia Genome Database (http://burkholderia.com) were performed. Based on these searches, none of the chokepoint enzymes in Table 3 had a mutant available; however, a *B. cenocepacia argG* mutant has attenuated virulence [178], and *recA* mutants have been identified in *B. cepacia* [179].

Additional metabolic enzymes, not identified as chokepoints in this study, and pathways critical for bacterial growth and survival have been mentioned with respect to target identification. These include anaplerotic pathways that turned on by limiting carbon sources [180], the glyxoalate shunt enzyme isocitrate lyase [16,181,182], involved in the metabolism of fatty acids [15,181], enoyl-ACP reductase (FabI) in the type II fatty acid biosynthesis pathway [183], and alanine racemase [23].

#### 4.4.2 Flux balance analysis

To gain an understanding of the metabolic processes in *B. pseudomallei* MSHR668 that are active under different environmental conditions, and to test the effect of deletion of each chokepoint enzyme on the growth of *B. pseudomallei in silico*, metabolic network models were constructed and FBA was performed. The first FBA model, of the unconstrained network in MSHR668, included all possible biomass compounds that could be produced, all nutrients that could be consumed, had no weights imposed on the biomass metabolites and no constraints imposed on the nutrients. [Supplementary material, S1_Final_unconstrained_model_inputs.pdf, S2_Final_unconstrained_model_solution.pdf]. This model likely represents the metabolic potential of *B. pseudomallei* in a soil or water environment where abundant carbon and nitrogen sources are available. To mimic the conditions that *B. pseudomallei* experiences in culture, a separate model was constructed that provided only the nutrients present in LB media plus glycerol [43]. Constraints were included on some of the nutrients in this model (ADP, Pi, proton and glycerol). [Supplementary material, S3_Final_LB_model_inputs.pdf, S4_Final_LB_model_solution.pdf]. A third model was attempted, to mimic infection conditions using only the nutrients present in the host cell cytosol. However, no comprehensive studies have identified a complete list of host cell nutrients that are available to *B. pseudomallei* during infection. Also, the specific carbon requirements of *B. pseudomallei* in either human macrophages or amoebae have not been determined. After trying to compile a list of nutrients that mimic the content of the host cytosol, from the literature and from gene expression studies of *B. pseudomallei* during infection, this model did not produce a solution so it was abandoned. However, the LB media model contained a similar set of nutrients to those used by the intracellular pathogen *L. pneumophila* [47,48,50],. so it may in fact be somewhat representative of infection conditions.

*In silico* knockout experiments were performed, where each chokepoint enzyme was knocked out, one at a time, to assess the effect on the total flux through the unconstrained and LB media models. Five chokepoint enzymes, when knocked out, had an effect on the models (Table 4). Specifically, knocking out BURPS668_3328 (tgt) and BURPS668_A2451 *(leuB),* eliminated the biomass flux in the unconstrained model, while knockout of BURPS668_2426 *(lpxD),* BURPS668_2433 (uppS), and BURPS668_3525 *(murG)* eliminated the biomass flux in the LB media model. Knockout of BURPS668_2433 (*uppS*), and BURPS668_3525 (*murG*) decreased the biomass flux in the unconstrained model, but did not eliminate it. The metabolic pathways that these enzymes belong to are shown in Figure 2.

**Figure 2.**
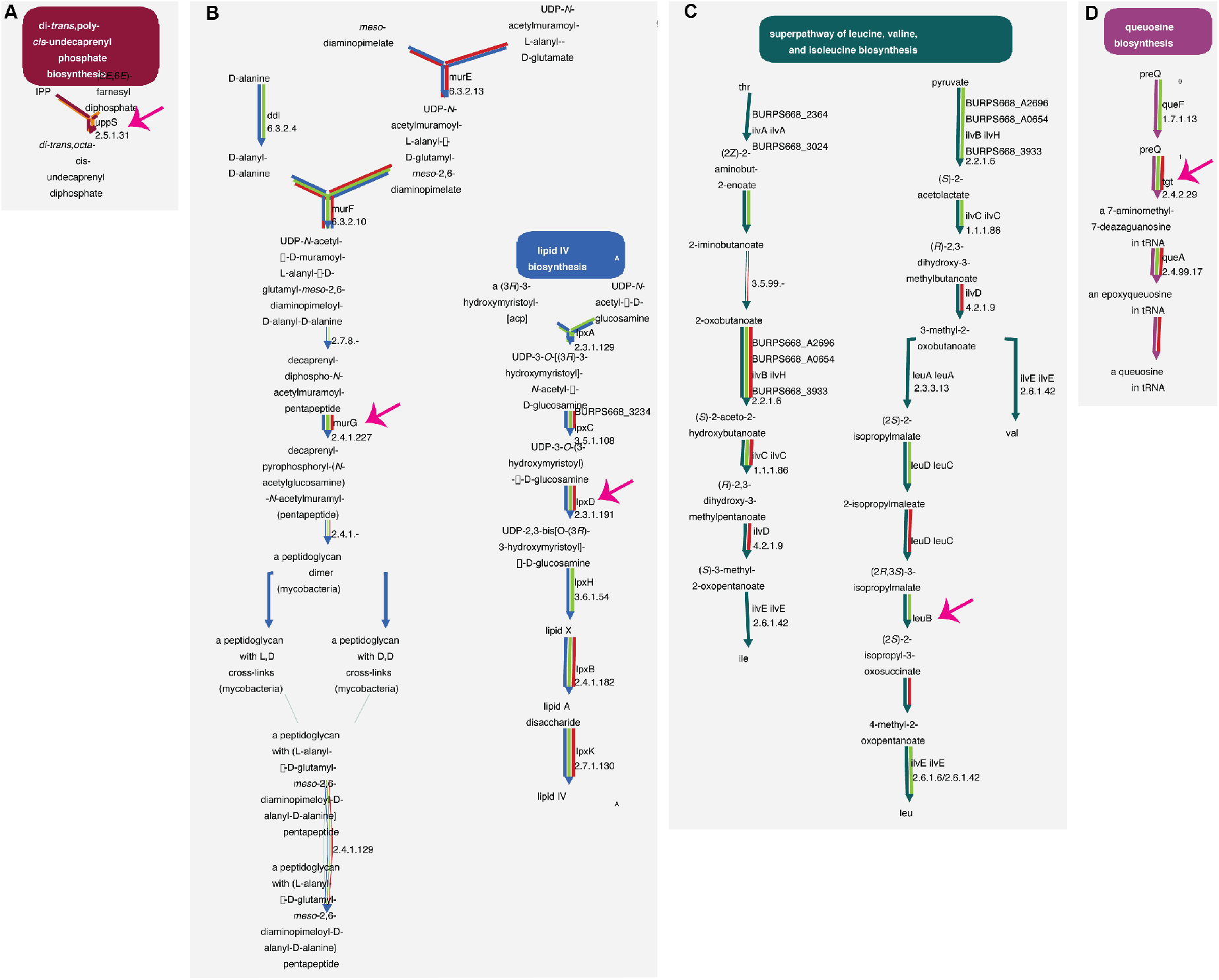
Metabolic pathways in *B. pseudomallei* MSHR668 that show reduced flux when chokepoint enzymes (indicated by pink arrows) are deleted. A. The mono-trans, poly-cis decaprenyl phosphate biosynthesis pathway that contains the chokepoint enzyme undecaprenyl diphosphate synthase (uppS). B. The two chokepoints UDP-N-acetylglucosamine--N-acetylmuramyl-(pentapeptide) pyrophosphoryl-decaprenol N-acetylglucosamine transferase *(murG)* and UDP-3-O-(R-3-hydroxymyristoyl)-glucosamine N-acyltransferase *(lpxD),* involved in peptidoglycan and lipid A biosynthesis, respectively. C. The chokepoint enzyme tRNA-guanine transglycosylase *(tgt),* involved in queosine biosynthesis. D. The 3-isopropylmalate dehydrogenase *(leuB)* chokepoint enzyme performs the third step in leucine biosynthesis. *In silico* deletion of UDP-3-O-(R-3-hydroxymyristoyl)-glucosamine N-acyltransferase (*lpxD*) reduced flux through the *B. pseudomallei* metabolic network in the LB media model, deletion of undecaprenyl diphosphate synthase *(uppS)* reduced flux through both unconstrained and LB media models, and deletion of tRNA-guanine transglycosylase *(tgt)* and 3-isopropylmalate dehydrogenase (*leuB*) reduced flux in the unconstrained model. These pathways were rendered by the Cellular Overview feature of Pathway Tools.

In terms of carbon sources, glucose was utilized as a nutrient in the unconstrained model of *B. pseudomallei* MSHR668. However, when glycerol was replaced by glucose in the LB media model, no FBA solution was found [data not shown]. This was a somewhat unexpected result, as *B. pseudomallei* can utilize glucose as a carbon source in culture [184]. One possible explanation for this result is that additional nutrients required for glucose utilization were missing from the input nutrients list. Cometabolism of more than one carbon substrate is a metabolic strategy employed by intracellular bacteria replicating inside host cells to provide carbon for energy and biosynthesis [59]. For example, *Listeria monocytogenes* can use both glycerol and lactate as carbon sources, [57,185–187]. It has been suggested that during infection by *Listeria,* the host cell may not contain enough glucose to activate bacterial PTS glucose transporters, so alternative carbon sources are important for survival and virulence of the pathogen [57]. *M. tuberculosis* also relies on glycerol and fatty acids as carbon sources in the macrophage environment [188] As the unconstrained *B. pseudomallei* model included a much longer list of nutrients than the LB model and also could use glucose as a nutrient, this could be the case. Another explanation for glucose not being utilized by the LB media model is that intracellular bacteria seem to prefer other substrates over glucose during infection [189], and glycerol may be a major carbon source for intracellular bacteria during infection [189,190]. This may also be the situation for *B. pseudomallei* inside host cells. Glycerol feeds into the second half of the glycolysis/gluconeogenesis pathway through its conversion to dihydroxyacetone phosphate [www.metacyc.org; [191]], bypassing the first four steps of glycolysis. We previously determined that the *B. pseudomallei* MSHR668 genome has the full set of genes to perform this conversion [37]. Two studies of *L. monocytogenes* infection support the idea that intracellular pathogens generally may use glycerol rather than glucose as a main carbon source while inside host cells. Transcription profiles of *L. monocytogenes* grown in mouse macrophages showed reduced expression of genes encoding some of the enzymes involved in glycolysis, in particular phosphoglucose isomerase (*pgi*), which converts glucose-6-phosphate into fructose-6-phosphate, and the five steps involved in the conversion of glyceraldehyde-3-phosphate to pyruvate [189]. Similar transcription profiles were seen during *L. monocytogenes* infection of Caco-2 epithelial cells [192]. Both studies showed increased expression of genes involved in the uptake and utilization of glycerol [189,192]: these genes were *glpF, glpK, glpD* and *dhaK.*

To date, no study has determined precisely which carbon substrates are utilized by *B. pseudomallei* during infection of host cells. In addition to glycerol, there is evidence that *B. pseudomallei* may utilize aromatic carbon compounds such as benzoate and phenylacetic acid as carbon sources for intracellular survival [53]. One study showed that in *B. pseudomallei* 1026b, glycolytic pathway and TCA cycle genes were down-regulated during infection of hamster [193], supporting the idea that *B. pseudomallei* may prefer carbon sources other than glucose while inside host cells. Other studies examined genes induced by hypoxia, which is a condition present in infected macrophages [194] and changes in *B. pseudomallei* gene expression during infection of rat lungs {van Schaik, 2008 #194.

Complicating the situation even more, the complete nutrient content of a representative mammalian host cell cytosol has not been determined yet, so a consensus set of nutrients present in the cytosol of different host cell types is still out of reach {Eisenreich, 2013 #40}. This is largely due to the challenges in designing appropriate infection models and robust analytical approaches to measure metabolic changes occurring in host cells during infection. Because of these limitations on both the pathogen and host sides, it is difficult to predict which carbon sources pathogens can use to grow inside host cells. While we don’t know the exact biochemical composition of a mammalian cell cytosol, we do know some details about mammalian cells in general. For example, the cytosol of a typical cell has low magnesium, sodium and calcium concentrations, and a high potassium concentration at neutral pH [195]. In addition, mammalian cells contain small amounts of amino acids, plus significant amounts of TCA cycle intermediates [196,197]. Once inside host cells, intracellular bacteria may stimulate host cell responses to produce needed nutrients [190]. However, host-pathogen interactions during infection are complicated, as some host defense responses are aimed at inhibiting pathogen survival and proliferation, for instance by decreasing metabolic activities that provide nutrients to the pathogen [180].

## 5. Conclusions

This work is the first to use genome scale metabolic modeling to address *B. pseudomallei* metabolism as a source of new drug targets. While identifying the nutrients available to *B. pseudomallei* inside host cells was difficult, the effort described here identified a set of twenty-six chokepoint enzyme drug targets; *in silico* deletion of five of these target enzymes reduced the total biomass flux through the *B. pseudomallei* metabolic network. While a genome-based approach like this can streamline the initial steps of antibacterial target identification, the true utility of this process will be demonstrated when the targets are experimentally verified by performing knockout experiments in culture, followed by efficacy testing of candidate drugs in culture and in animal models of infection.

## Supplementary Materials

The following are available online at www.mdpi.com/link, S1_Final_unconstrained_model_inputs.pdf,S2_Final_unconstrained_model_solution.pdf, S3_Final_LB_model_inputs.pdf, S4_Final_LB_model_solution.pdf, S5_Nutrients_infection_model.pdf

## Acknowledgments

This project was supported by the Defense Threat Reduction Agency (JSTO-CBD Proposal # CBCALL12-IS1-1-0283).

